# A mechanical wave travels along a genetic guide to drive the formation of an epithelial furrow

**DOI:** 10.1101/2022.12.08.518365

**Authors:** Anna Popkova, Sophie Pagnotta, Matteo Rauzi

## Abstract

Epithelial furrowing is a morphogenetic process that is pivotal during embryo gastrulation, neurulation and the shaping of the animal body. A furrow often results from a fold that propagates along a line. How fold formation and propagation are initiated, driven and controlled is still poorly understood. To shed new light on this fundamental morphogenetic process, we study the formation of the cephalic furrow: a fold that runs along the dorsal-ventral axis of the embryo during early *Drosophila* gastrulation and the developmental role of which is still unknown. Here, we provide evidence of its function and show that the cephalic furrow is initiated by two groups of cells located on the left and right lateral sides of the embryo. These cellular clusters work as a pacemaker triggering a bi-directional morphogenetic wave powered by actomyosin contractions and sustained by *de novo* medial apex-to-apex cell adhesion. The Cartesian position of the pacemakers is under the cross-control of the embryo anterior-posterior and dorsal-ventral gene patterning systems. Thus, furrow initiation and propagation are driven by a mechanical trigger wave that travels under the control of a multidimensional genetic guide.

## Introduction

Epithelial furrowing is a remarkable folding process that involves the coordinated bending of a strip of cell tissue. This morphological transformation is generally known to be key during the formation of the neural tube in verebrates^1^ but it has also been shown to play a pivotal role during embryo gastrulation^2,3^ and body or organ shaping^4-6^. Epithelial furrowing often takes place in a stereotypic fashion: a fold forms at one position of a strip of tissue and eventually propagates along the tissue plane following a linear trajectory^7-10^. How fold formation and propagation are controlled and driven is still poorly understood. To tackle this, we investigate the process of cephalic furrow (CF) formation during the early gastrulation of the *Drosophila* embryo.

CF formation is among the first morphogenetic changes characterizing the onset of *Drosophila* gastrulation, therefore marking the blastula-to-gastrula transition. From this perspective, it is an ideal model process to study epithelial furrowing because the initial starting conditions are simple with cells being in a steady state. The genetic control of the CF is quite well deciphered. Overall, it is known that the CF position along the embryo anterior-posterior (AP) axis and its formation are under the control of the AP gene patterning system^11-14^. The CF runs along a line parallel to the dorsal-ventral (DV) axis on both lateral left and right sides of the embryo and it is positioned roughly at one third of the embryo length from the anterior pole separating the head from the trunk region. The folding process can be divided into two phases: a first phase named “apical-basal shortening” in which a cell (dubbed initiator cell or IC) reduces its apical-basal height forming a groove under the control of Rho^15^, and a second phase named “rolling over” in which cells surrounding the IC sequentially move into the groove^14,16^. While a recent study has focused on better deciphering the gene expression and morphogenetic noise and their impact on CF linearity, how the CF forms and propagates is still not clear. Intriguingly, the CF is a reversible fold that eventually unbuckles. The developmental role of this transient furrow or of its folding and unfolding is yet unknown. By using two-photon optogenetics coupled with multi-view light sheet microscopy, laser-based manipulation, across-scales analysis and multidimensional image processing, here we seek to unravel the fundamental mechanisms controlling and driving fold formation and propagation resulting in an epithelial furrow.

## Results

### Tissue furrowing is driven by intrinsic active forces

Tissue folding can be initiated by extrinsic or intrinsic mechanical forces. In-plane movement of the tissue can drive epithelial folding^17-19^ or can contribute to tissue involution^20-24^. We thus wondered if the initiation of the CF is the result of tectonic buckling driven by extrinsic in-plane compressive stresses between the trunk and the head tissues moving against each other. To address this question, we monitored cell apices position over time along a coronal section during CF formation (Fig. 1a). If cells at different positions away from the furrowing point move together towards the IC, then the tectonic buckling hypothesis is not to be discarded. Our analysis shows that the IC moves before and faster than the other cells positioned further away from the furrowing point (Fig. 1b). This evidence does not support the tectonic buckling hypothesis and favors an active IC mechanism. To directly rule out the hypothesis that distant movement of trunk and head tissues towards the furrowing zone initiate CF formation, we took advantage of the cauterizing capabilities of the infrared femtosecond laser (IR fs). Laser cauterization, if performed at the interface between the embryo vitelline membrane and the blastoderm epithelium, has been shown to drive the formation of immobile boundaries that act as physical barriers^20,25-27^. We generated immobile boundaries around the furrowing zone before the onset of CF formation in order to ultimately protect the CF area from in-plane compressive stresses (Fig. 1c). After tissue cauterization, the CF still forms without a noticeable delay (Supplementary Fig. 1a and b) and the tissue tears apart from the cauterized zone (Fig. 1c yellow arrowhead and Supplementary movie 1). This evidence definitely rules out the tectonic buckling hypothesis and demonstrates that active forces are generated within the folding region to drive CF formation.

**Figure 1.**
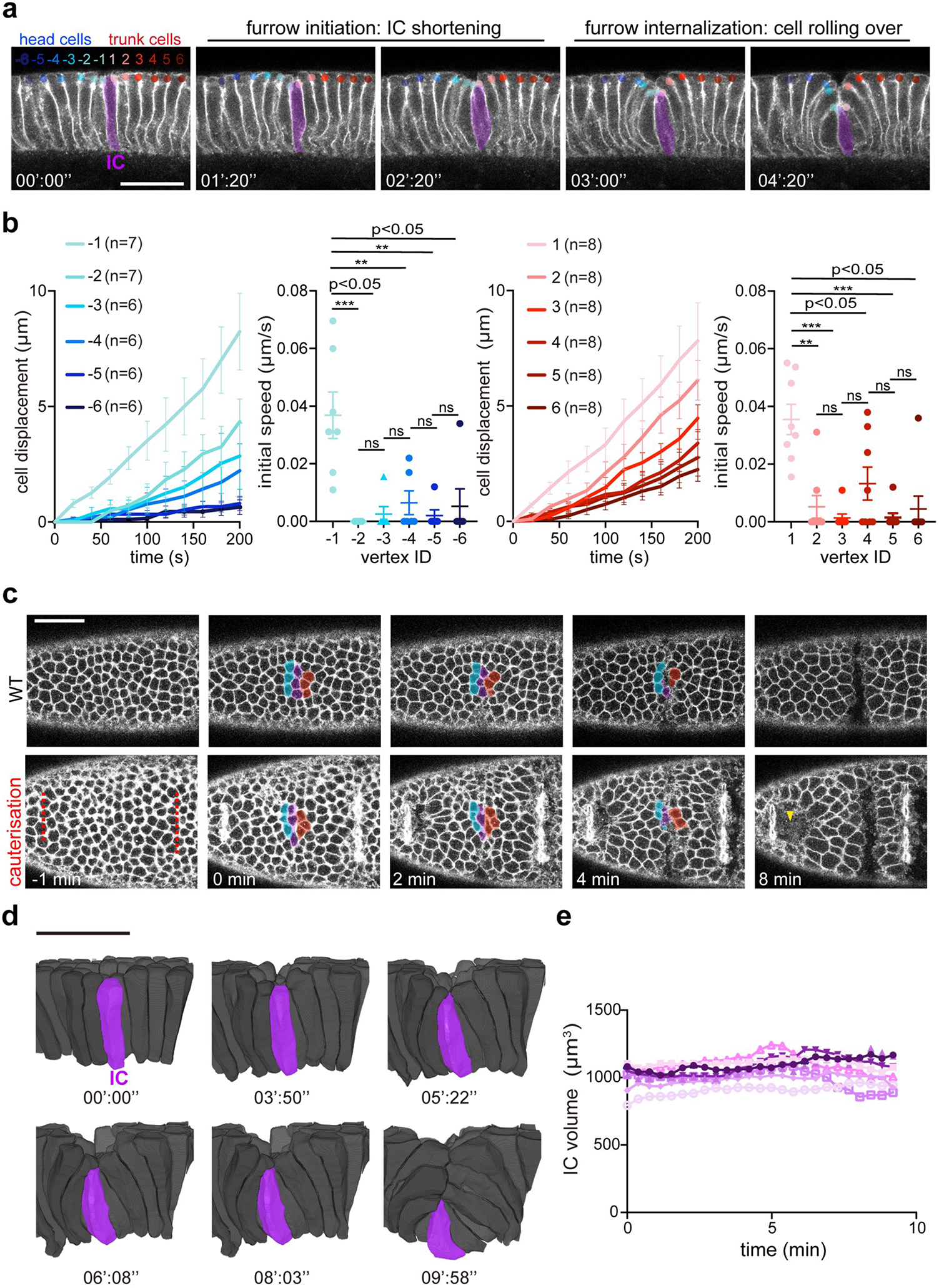
CF formation is initiated by intrinsic forces. **a** Mid-coronal time-lapse imaging of CF formation in a membrane labeled *klar-* embryo to increase epithelial transparency for deep tissue imaging. The IC is in purple. t = 0, onset of CF formation. Colored circles label apical extremities of the IC (−1 and 1), head cells (−1 to -6) and trunk cells (1 to 6). n indicates the number of CFs. **b** Cell apical extremity displacement as a function of time. Error bars: ±SEM. n indicate number of CFs. **c** Time-lapse en face view of control (top) and cauterized (bottom) embryos. ICs in purple, neighbor head cells in blue and neighbor trunk cells in red. **d** Rendering of 3D cell segmentation time-lapse during CF formation in a membrane labeled *klar*-embryo. **e** Plot of IC volume over time. n = 8 ICs, 3 embryos. Scale bars 30 μm.

Changes in cell volume have been shown to be an active mechanism that drives tissue morphogenesis^28^. We therefore tested if changes in cell volume are responsible for generating intrinsic active forces that direct CF initiation resulting in the shortening of the IC. To test for this, we imaged CF formation with multi-view light sheet microscopy and implemented 4D data registration and fusion providing 200 nm isotropic resolution images for accurate cell segmentation (Fig. 1d and Supplementary movie 2). 4D cell morphometric analysis shows that the IC volume does not undergo major variations over time (Fig. 1e). Finally, CF initiation is not induced by tectonic buckling but is driven by active mechanical forces generated within the CF tissue that do not involve IC volume change.

### RhoGEF2, Dp114RhoGEF, Rho and ROCK are necessary for furrow formation

Active mechanical forces are often driven by the actomyosin cytoskeleton under the control of the Rho pathway. We thus thoroughly investigated the function of this pathway in controlling and driving CF formation. RhoGEF2 and Dp114RhoGEF are two Rho guanine nucleotide exchange factors activating the Rho pathway that have been shown to be key players during lateral tissue morphogenesis in the early *Drosophila* embryo^29^. To test the role of these two RhoGEF factors, we downregulated RhoGEF2 and Dp114RhoGEF by RNA inhibition. Both RhoGEF2 and Dp114RhoGEF RNAi show CF formation delay or even failure with RhoGEF2 RNAi showing a stronger failure phenotype (Fig. 2 a, b and Supplementary Fig. 2a). We then tested the epistatic interaction of the two RhoGEFs. In RhoGEF2-Dp114RhoGEF double RNAi, CF formation is further delayed and more than half of the embryos showed CF failure (Fig. 2b and Supplementary Fig. 2a). This shows that RhoGEF2 and Dp114RhoGEF act together to ensure CF formation in a timely fashion.

**Figure 2.**
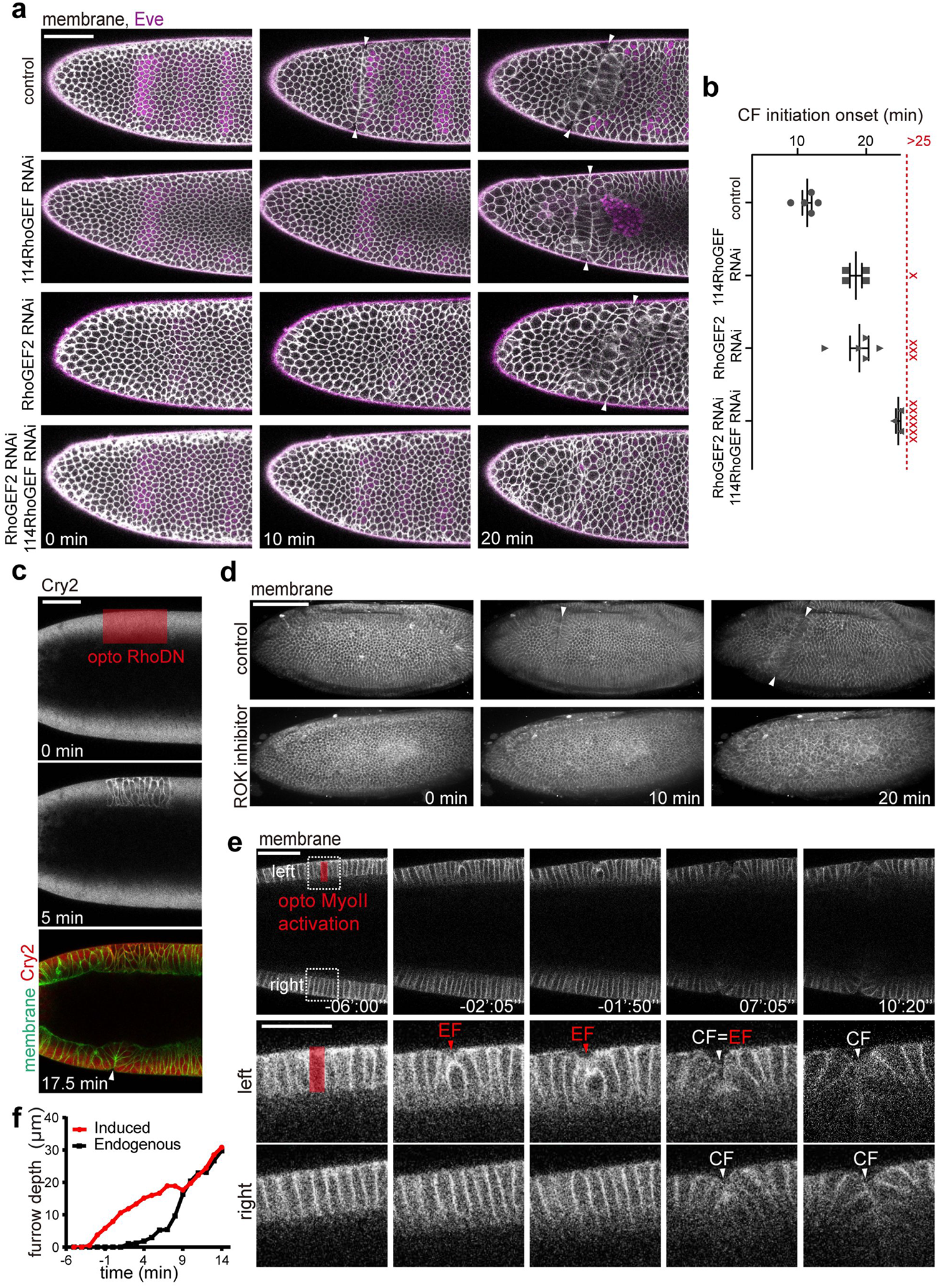
Dissecting the Rho pathway in CF formation. **a** Time-lapse images of control and RhoGEF RNAi embryos. t = 0 indicates 30 μm of cellularization front. **b** Plot showing the CF initiation onset in the indicated genetic backgrounds. One data point per embryo. Error bars: ±SEM. **c** Time-lapse images of an embryo co-expressing CIBN::pmGFP (green) and mCherry::Cry2::RhoDN (red). Before illumination, Cry2::RhoDN is cytosolic. After two-photon laser activation, Cry2::RhoDN is cortically recruited in the region of activation. t = 0, just before two-photon activation. **d** Time-lapse images of embryos injected with water and ROCK-inhibitor. **e** Time-lapse images showing MyoII activation in a presumptive IC on the left side of the embryo prior to CF formation. Top panel: zoom-out mid-coronal view. Bottom panels: zoom-in views of the photo-activated (left) and control (right) regions. EF, ectopic fold. Red rectangle denotes the photo-activated region. **f** Furrow depth over time in photo-activated (red) and control (black) folds as shown in **e**. Scale bar 50 μm (**a, c, e** top panel), 100 μm (**d**) and 30 μm (**e**, bottom panels).

A previous study has provided evidence that Rho1 is necessary for CF formation^15^. Since the CF initiates its formation on the two lateral sides of the embryo at about the same time (Supplementary Fig. 2b)^15^, we now aimed to validate the role of Rho and directly test if the formation of the left and right folds are functionally coupled from one another. To that end, we performed two-photon activation of a Rho dominant negative (RhoDN) optogenetic construct on one single side of the embryo while monitoring both left and right regions. The formation of the CF was inhibited on the activated side only (Fig. 2c) demonstrating that the formation of the left and right folds can be uncoupled and validating previous results^30^. Rho-associated protein kinase (ROCK) is known to act downstream of Rho to activate actomyosin contractility. We therefore injected a ROCK inhibitor (Y-27632) at the end of cellularization (before CF initiation) to test directly the role of this kinase in driving CF formation. After Y-27632 injection the CF failed to form showing that ROCK is an essential kinase to drive this process (Fig. 2d and Supplementary movie 3).

CF formation is initiated by one or two ICs that undergo apical-basal shortening^15,31^. We thus hypothesized that activation of Rho in one or two cells along the lateral side of the embryo is sufficient to initiate cell apical-basal shortening. To test for this, we implemented two-photon optogenetic RhoGEF2 to activate cell cortical actomyosin contractility^32^ in one or two cells near the CF area. The activated cells underwent apical-basal cell shortening forming an ectopic groove (Supplementary Fig. 2c). The groove eventually unfolded when CF formation initiated (Supplementary Fig. 2c, last panel and Supplementary movie 4), probably as a result of in-plane traction forces generated by the CF (Fig. 1c). We then activated myosin II (MyoII) in an IC to test if we could trigger CF formation. We activated an IC on the left side while monitoring both left and right tissues. After activation, the IC shortened along the apical-basal axis forming an ectopic groove as in the previous experiment. Nevertheless, the groove, after reaching ∼20 μm in depth, stalled (Fig. 2e and f). The right fold eventually started to form (Fig. 2e and f at time point 0) and, after reaching a depth similar to the one of the ectopic fold, both right and left folds moved inside synchronously (Fig. 2f and Supplementary movie 5). This shows that, while cell scale activation of the Rho pathway is sufficient to drive cell apical-basal shortening, CF formation cannot be triggered by ectopically driving IC shortening. Therefore, while formation of left and right folds are uncoupled from one another (i.e., the fold on one side can form while the fold on the other side is inhibited), they both depend on a higher order program controlling and synchronizing their formation.

### Cell lateral contraction is effective to initiate fold formation

Our experimental trials have shown that cortical recruitment of RhoGEF2 (controlling the activation of actomyosin contractions) in a single cell is sufficient to drive cell apical-basal shortening (the first step in CF formation). To better decipher the sub-cellular distribution of MyoII during CF formation, we performed enhanced resolved confocal imaging at high magnification to quantitatively monitor MyoII dynamic recruitment on different cell sides. MyoII is enriched basally already several tens of minutes before CF onset during a process named cellularization (the process of cell formation that starts before and continues during the beginning of CF formation). At the onset of CF formation, MyoII accumulates first apically in the CF zone and then laterally^15^ along the sides of the IC (Fig. 3a and Supplementary movie 6). Therefore, three pools of MyoII can be identified in the CF area during CF formation: cell apical, lateral and basal pools.

**Figure 3.**
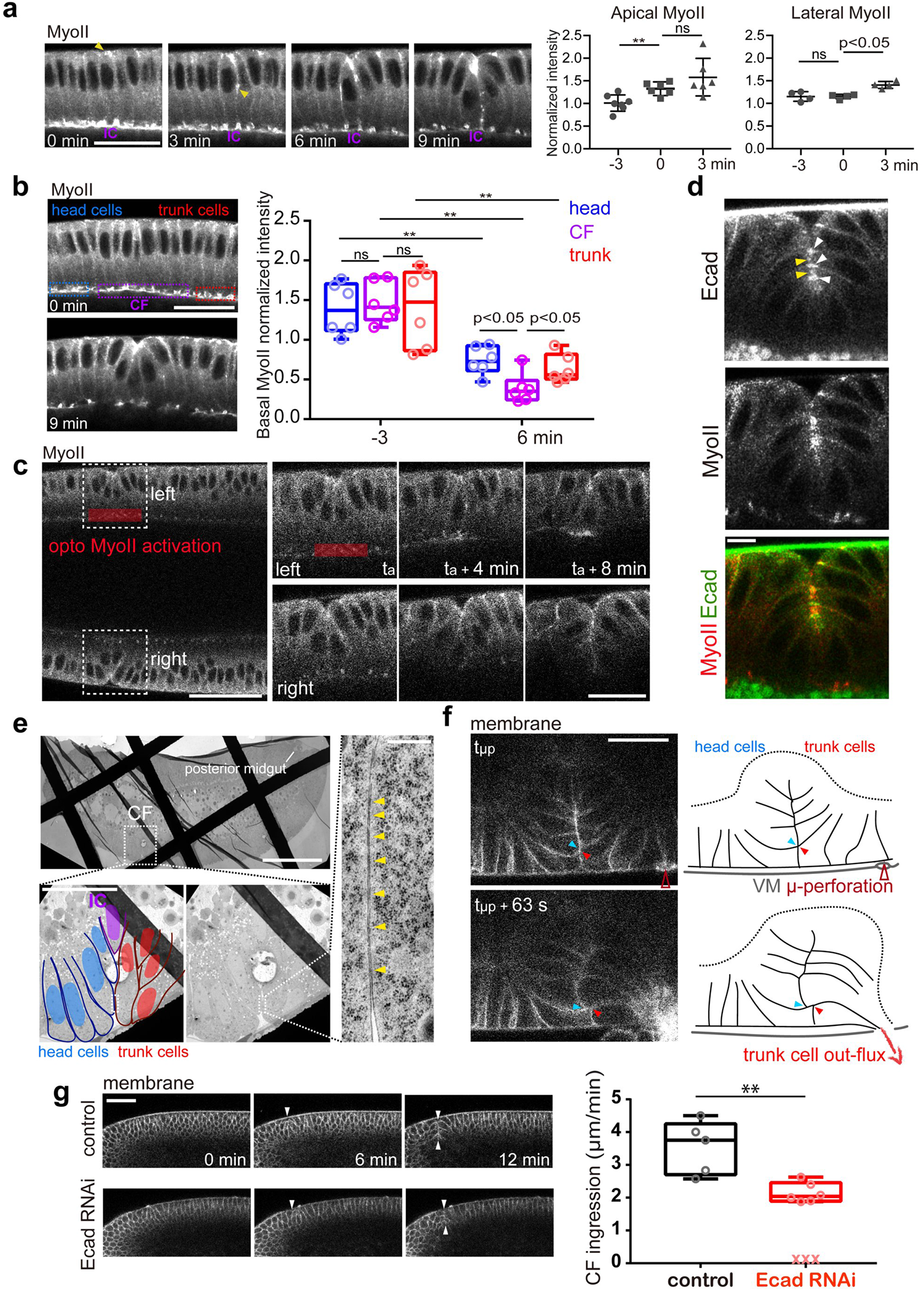
Dissecting the contractile machinery driving CF formation. **a** Mid-coronal time-lapse images showing MyoII distribution during IC shortening (left). t = 0 indicates the onset of CF formation. Plot of MyoII intensity on the apical and lateral sides of ICs. Error bars: ±SD. **b** Basal MyoII distribution before and during CF formation. Box plot indicates median with max. and min. values. **c** Basal MyoII activation. Red rectangle indicates the activation zone. Zoom-in time-lapse images of the left (activated) and right (control) CF folds (dashed frames). **d** High magnification imaging of the CF. White and yellow arrow heads indicate apical-lateral and apical-medial E-cadherin clusters, respectively. **e** Electron micrograph of the embryo during CF formation. Top is a Zoom-out, bottom-left is a zoom-in with head cells in blue, trunk cells in red and the IC in purple. Right panel shows high magnification of the contact zone between a head cell and a trunk cell. Arrowheads indicate electron dense regions between the two facing cell cortexes. **f** IR fs laser perforation (brown arrowhead) of the vitelline membrane in the proximity of CF causing tissue out-flux. Blue and red arrowheads indicate sliding of a pair of head and trunk cells. n = 4 folds from 4 embryos. **g** Furrow formation in a control (water-injected) and an E-cadherin RNAi-injected embryo. The plot indicates the speed of CF ingression in the two cases. A CF per data point. Scale bar 30 μm (**a, b**), 50 μm (**c** left and **g**), 10 μm (**d**), 100 μm (**e** top left), 25 μm (**c** right, **e** bottom left and **f**) and 0.5 μm (**e** right). A CF per data point.

A cell in a monolayer epithelium has three distinct sides: the apical, basal and lateral sides that define the boundary between the cell and the outside, the cell and the inside and between cell neighbors, respectively. Therefore, we now investigated which side of a cell would mostly contribute to cell apical-basal shortening if subjected to contractile forces by implementing two-photon sub-cellular activation of optogenetic RhoGEF2. Activation of the apical and basal sides resulted in mild and no apical-basal shortening, respectively (Supplementary Fig. 3a and c). Activation of the lateral side, however, resulted in strong shortening of cells along the apical-basal side (Supplementary Fig. 3b). In order to test if MyoII activation on one side is sufficient to drive cell apical-basal shortening, we photo-activated RhoGEF2 on one single lateral interface. Remarkably, this results in apical-basal shortening and an eventual sharp bent in the tissue (Supplementary Fig. 3d). Overall, this shows that lateral contractions are effective in driving cell apical-basal shortening therefore corroborating the idea that it is mainly lateral contraction of the IC that initiates CF formation, as previously suggested^15^.

### Tissue involution is driven by basal loss of MyoII and is sustained by medial apex-to-apex cell adhesion

We have shown that MyoII activation at the basal side of CF cells does not contribute to cell shortening and CF initiation. During CF formation, MyoII at the basal side of the bending zone is more strongly downregulated compared to neighboring regions (Fig. 3b). We therefore hypothesized that MyoII downregulation is key for CF formation (similar to during the formation of the ventral fold^33^). To test for this, we used two-photon optogenetics to activate MyoII at the basal side of CF cells. Remarkably, CF rolling over was compromised (Fig. 3c), supporting the idea that enhanced basal MyoII downregulation in the bending region is functional for CF rolling over. We then questioned whether basal downregulation of actomyosin contraction would be sufficient to drive CF rolling over. To test for this, we used IR fs laser dissection of the actomyosin network on the cell basal side. After ablation along a line, a furrow forms, leading to partial tissue involution (Supplementary Fig. 4a). This corroborates the notion that downregulation of basal MyoII and basal tension is necessary and sufficient to drive CF involution.

During rolling over, opposing trunk and head cell apices are juxtaposed with another^31^. We therefore hypothesized that trunk and head cells come in apical contact during CF rolling over. To test for this, we monitored E-cadherin distribution in CF cells: E-cadherin is initially localized in the sub-apical zone of the cell and eventually relocalized apically^31^. When the apex of trunk cells approaches the apex of head cells, E-cadherin clusters are focused on the apical lateral side of both opposing cells. Since opposing cells can face each other, apical lateral E-cadherin appears to form clusters across the trunk and head cell apices (Fig. 3d, white arrow heads) as if initially distant trunk and head cells would eventually join together by forming contacts. During CF rolling over, E-cadherin not only forms matching clusters along the apical lateral side but also forms clusters in the apical medial zone (Fig. 3d, yellow arrow heads and Supplementary Fig. 4b), leading to the hypothesis that the process of CF formation results in the formation of new contacts between opposing trunk and head cells (i.e., the medial apex-to-apex adhesion hypothesis). To test this hypothesis, we performed electron microscopy (EM) imaging during CF rolling over to optically resolve any spot adherens junctions between facing trunk and head cells. EM images show facing apical membranes at a distance of 10-20 nm decorated with discontinuous electron dense regions: this is a characteristic EM pattern formed by cell-cell spot adherens junctions^34^ (Fig. 3e, between a trunk cell – blue – and a head cell – red). We then aimed to test directly the medial apex-to-apex adhesion hypothesis. To that end, we developed a new technique to generate in-plane ectopic traction forces that would pull the trunk tissue away from the head tissue eventually driving CF unfolding. If trunk and head cells within the CF were strongly adhering one another at their apices, trunk and head cells would remain in close proximity and the CF would be protected from unfolding. In order to generate in-plane pulling forces we induced μ-perforation of the vitelline membrane by exploiting the multi-photon surgical effect of the IR fs laser. Since the embryo inside the vitelline membrane is under pressure, μ-perforation of the vitelline membrane results in an out-flux of cellular material generating pulling forces. After performing a μ-perforation on the trunk side in proximity to the folded region, the CF remains intact without unfolding and opposing trunk and head cells remain in close proximity (Fig. 3f). Remarkably, during cell out-flux, trunk cells engaged in the CF can eventually drag their head cell partner while pulled by ectopic traction forces (Fig. 3f, red and blue arrowheads and Supplementary movie 7). This directly demonstrates that head and trunk cells adhere to one another during CF formation. In order to test the role of cell-cell adhesion in CF formation, we injected *shotgun* RNAi to downregulate E-cadherin levels. *shotgun* RNAi embryos show delayed or compromised CF formation compared to water-injected control embryos (Fig. 3g). Therefore, we can conclude that CF formation involves the creation of adherens junctions between medial apices of originally distant trunk and head cells and that E-cadherin based adhesion is functional for CF formation.

### A trigger wave propagates under the control of a multidimensional genetic guide to drive furrow formation

After unravelling the mechanisms responsible for CF formation at the cellular and sub-cellular scales, we now aimed to dissect this process at the tissue and embryo scales. The CF develops on both right and left sides of the embryo and along a strongly curved surface (i.e., along the embryo DV axis). Therefore, it is challenging to image the entire folding process over time and conventional fluorescence confocal microscopy fails to provide a global view of CF formation. To overcome this challenge, we implemented multi-view light sheet microscopy to image the entire embryo over time with sub-micron isotropic pixel resolution and performed cylindrical projections of the membrane-tagged blastoderm after 3D isotropic digital reconstruction^20,35^. Projections are effective to monitor the formation of the CF from the ventral to the dorsal region on both left and right side simultaneously (Fig. 4a). Our data show that the left and right segments of the CF start forming on the two lateral sides of the embryo (in the same region where nuclei first move towards the basal side^15^) and eventually propagate towards the ventral region (intersecting the ventral fold) and towards the dorsal region merging together (Fig. 4a).

**Figure 4.**
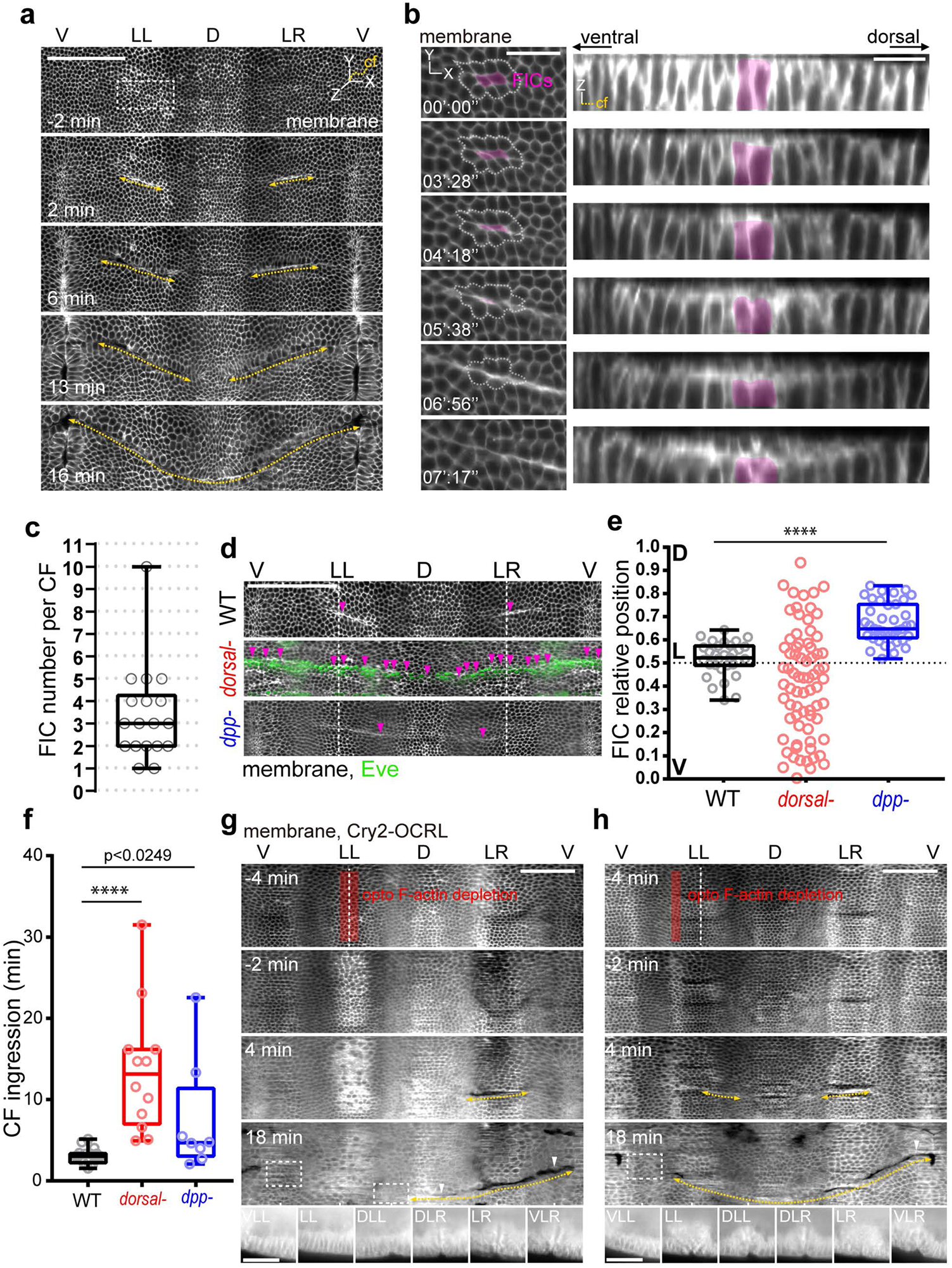
The CF is formed by a trigger wave initiated by FICs. **a** Cylindrical projection of the embryo anterior side. Ventral (V), lateral left (LL), dorsal (D), lateral right (LR). The yellow dashed line indicates CF propagation. **b** Time-lapse images showing first initiator cells (FICs, purple) in a en face view (left) and along a section following the CF (right). **c** Number of FICs per CF. n = 18 CFs. **d** and **e** FIC position (purple arrowheads in **d**) in wild type, *dorsal*-mother and *dpp*-embryos. **f** Time span between FIC apical narrowing and CF ingression in wild type, *dorsal*-mother and *dpp*-embryos. **g** Localized disruption of the actin cytoskeleton (red rectangle) with optogenetics on the lateral left side of the embryo where FICs are positioned. The yellow dashed line indicates CF propagation and dashed white rectangles indicate regions of CF inhibition. At the bottom, ventral lateral left (VLL), lateral left (LL), dorsal lateral left (DLL), dorsal lateral right (DLR), lateral right (LR), ventral lateral right (VLR) sections of the CF at 18 min. **h** As in **g** with optogenetic activation on the ventral lateral side. Scale bar 100 μm (**a, d, g** and **h**), 50 μm (bottom panels in **g** and **h**) and 20 μm (**b**).

A previous study has shown that during mesoderm invagination, the ventral fold propagates in the absence of a cell MyoII-constriction wave^7^. This propagation phenomenon results from long-range mechanical tissue interactions powered by synchronous MyoII recruitment and concerted constriction of the mesoderm cells. To test if a similar mechanical process was driving CF formation, we monitored CF cells along cylindrical projections. Remarkably, CF cells do not undergo apical-basal shortening synchronously: a cluster of two to three ICs, in average, first undergo cell shape changes (Fig. 4b and c) which initiate a morphogenetic wave. We thus named these cells *first initiator cells* (FICs). In the CF, cell shape change propagation is mirrored by a MyoII wave that also initiates on the lateral sides of the embryo and eventually spreads towards the ventral and dorsal regions along the furrow line (Supplementary Fig. 5a and Supplementary movie 8). This shows that the mechanism underlying the morphogenetic propagation of the ventral fold and the CF is different. How is FIC position determined? Since FICs are located at a specific DV position along the CF, a DV signal is a priori necessary to provide DV spatial information to the cells. We therefore screened for DV patterning genes that could potentially carry the necessary spatial information. Remarkably, *dorsal*^36,37^ and *dpp*^38,39^ gene mutations showed a FIC phenotype. For embryos from females carrying a *dorsal* homozygous mutation (*dorsal*-), FICs are no longer spatially constrained but are spread along the DV axis, while for embryos carrying a *dpp* mutation (*dpp*-), FICs are shifted to a more dorsal position compared to that in wild type embryos (Fig. 4d and e and Supplementary movie 9, 10). While in *dpp*-embryos the CF does not show a major alteration in time or dynamics, in embryos from *dorsal*-mothers the CF is strongly delayed (Fig. 4f) and forms simultaneously at different DV positions so fails to propagate (Supplementary Movie 11). This acute phenotype could be explained by the fact that embryos from *dorsal*-females lack the MyoII propagation wave (Supplementary Fig. 5b and Supplementary movie 12) similar to AP patterning mutant embryos (e.g., embryos from *bnt*-mothers, Supplementary Fig. 5c and Supplementary movie 13) in which CF formation is permanently abolished^11,14^. Other processes at a later time (e.g., residual posterior midgut movements or mitotic cell divisions - Supplementary Fig. 5d) still occurring in *dorsal*-embryos, may eventually play a role in rescuing CF formation in such a compromised genetic background.

Morphogenetic propagation in an epithelium can be driven by waves of different nature. The two main wave types are trigger and phase waves. While the former relies on cell-cell interaction and occurs when an excitable unit kicks the neighbor excitable unit and so on^40^, the latter is interaction independent and results from a cell-to-cell phase shift^41^. Therefore, a corollary is that a discontinuity in cell-cell interaction can halt a trigger wave but not a phase wave. In order to directly test if CF formation is a morphogenetic trigger or phase wave, we aimed to ectopically generate a physical discontinuity with spatial and temporal precision along the CF line. To that end, we designed an experimental trial based on two-photon optogenetics to perturb the F-actin cytoskeleton with spatio-temporal precision^42^ coupled with multi-view light sheet microscopy^25^ for *in toto* imaging and cylindrical image projection. In this experiment, we performed photo-activation on one single lateral side (e.g., the left side) of the embryo in order to use the other side (i.e., the right side) as an internal control. When photo-activating a sub-region located on the most lateral left side (where FICs are usually located - Fig. 4g, red), the CF is entirely abolished on the left-activated side and preserved on the right-control side (Fig. 4g). This shows that FICs function as a pacemaker initiating the morphogenetic wave. Interestingly, at a later time when the posterior midgut moves along the dorsal side of the embryo towards the anterior pole, the CF can eventually still form on the dorsal-left side but not on the ventral-left side (see Supplementary movie 14). This shows that CF formation on the dorsal side is more robust than on the ventral side, probably being dorsally driven by multiple redundant mechanisms. We then activated a sub-region, this time located in the ventro-lateral left side of the embryo (Fig. 5h, red), in order to preserve FIC cytoskeleton integrity and monitor the propagation of the morphogenetic wave. This time, the morphogenetic wave is also initiated on the left side of the embryo (eventually merging dorsally with the unperturbed right furrow) but it is halted on the ventral-lateral zone that was previously photo-activated (Fig. 5h, dashed box and Supplementary movie 15). This directly demonstrates that CF formation is a trigger wave initiated by a patch of contracting cells (i.e., the FICs) that act as a pacemaker stimulating the surrounding excitable tissue strip.

**Figure 5.**
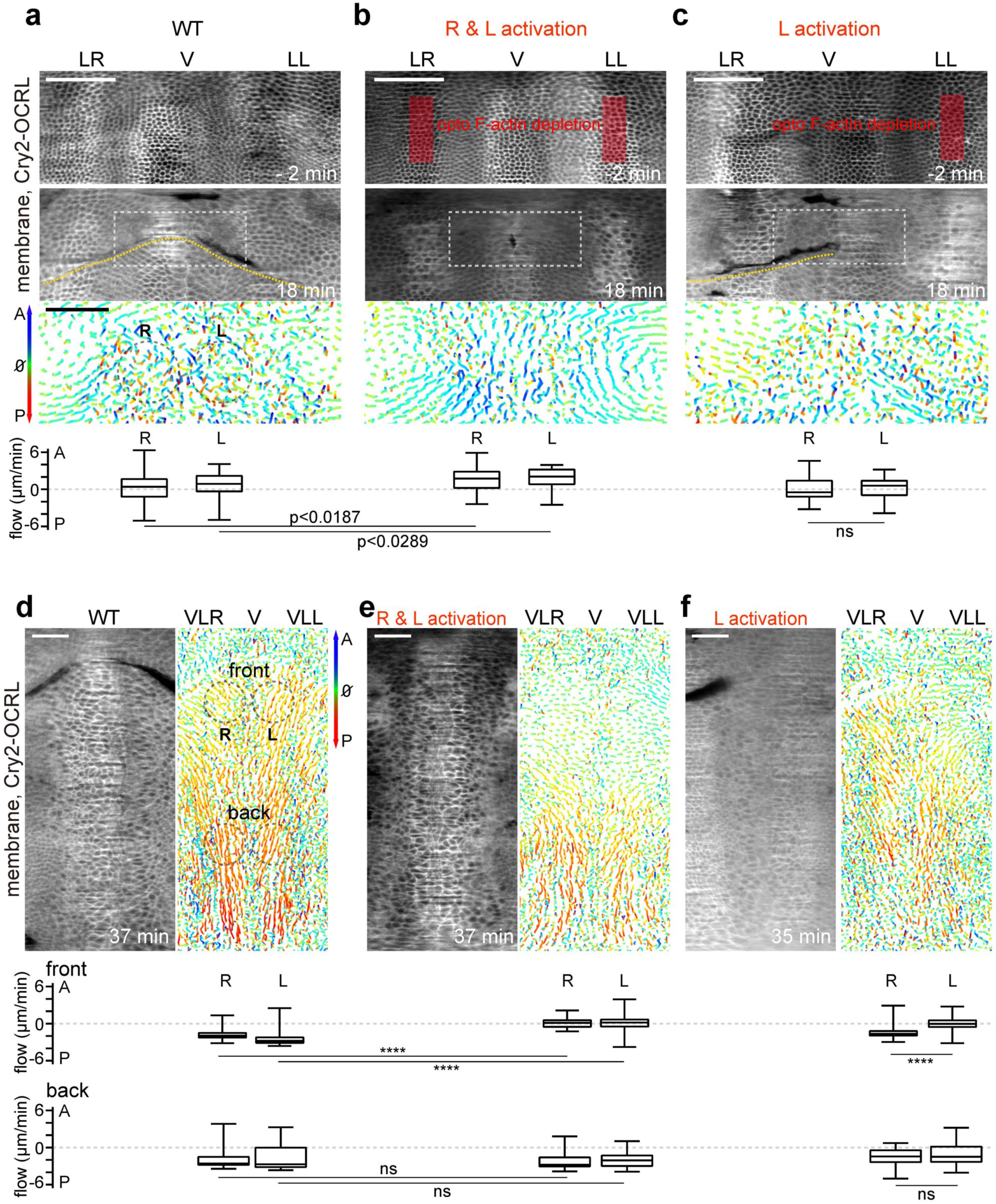
The CF works as a morphogenetic barrier and an epithelial reservoir. **a** Cylindrical projection of the embryo anterior side. Lateral right (LR), ventral (V), lateral left (LL). The yellow dashed line indicates the propagation of the CF. The third panel shows the cell flow during CF formation within the dashed white rectangle. Cell forward (towards the anterior - A) and backward (towards the posterior - P) flow is measure for the right (R) and left (L) sides. **b** as in **a** in the absence of both lateral and right CF folds inhibited with optogenetics (red rectangle). **c** as in **b** in the absence of the left CF fold. **d** Cell flow during ventral lateral CF unfolding. Ventral lateral right (VLR), ventral (V), ventral lateral left (VLL). Scale bar 100 μm (**a, b, c**) and 50 μm (**d, e, f**).

### The dual function of the CF: a morphogenetic barrier inhibiting forward cell flow and an epithelial reservoir powering backward cell flow

During embryogenesis, tissue folding is often an irreversible process that is essential to translocate cells to the interior of the embryo where inner organs are eventually formed. CF formation is instead a reversible process: at later stages of embryo development, the CF unfolds. What is the function of the CF and how does sequential folding and unfolding contribute to the development of the embryo? In 1970’s, F. R. Turner and A. P. Mahowald speculated that the CF could function as a temporary epithelial “storage”^43^, while in a recent computational study, researchers have hypothesized that the CF could work as a morphogenetic barrier^44^. In order to test these hypotheses, we developed an experimental design to abolish the CF on both sides of the embryo and monitor cell flow on the embryo ventral zone where the CF is permanently abolished. To that end, we performed ambilateral two-photon optogenetics coupled with multi-view light sheet microscopy, cylindrical image projection and cell flow analysis (see Methods). Our experimental trials show that in an embryo deprived of CF, a stronger forward directed flow (towards the anterior) is generated compared to control conditions (Fig. 5a and b). This shows that the CF can work as a morphogenetic barrier, reducing forward flow and thereby isolating the head region of the embryo. During the slow phase of germband extension, the CF unfolds on the ventral side (see Supplementary movie 16)^45^. During CF ventral unfolding, a backward directed flow (towards the posterior) is generated both on the front and back trunk region of the embryo (Fig. 5d). Remarkably, in embryos deprived of CF, the backward flow is maintained on the back but is absent on the front (Figure 5e and Supplementary movie 17). We then inhibited the CF only on one side of the embryo. While the forward flow did not show significant difference between right and left sides, the backward flow at the front on the activated side was significantly less than on the control side (Fig. 5c and f and Supplementary movie 18). Finally, this shows that the CF also works as a tissue reservoir that can eventually power cellular flow during unfolding.

## DISCUSSION

Epithelial furrowing plays a paramount role during embryo development. The mechanisms and mechanics controlling and driving this remarkable folding process are still poorly understood. The furrowing of tissues often results from the formation of a fold that propagates along a line. Understanding how furrow propagation is triggered and powered is therefore key to unveil the fundamental principles underlying epithelial furrowing. As a paradigm, we study the formation and propagation of the CF during early *Drosophila* gastrulation: a furrow that separates the head from the trunk region of the embryo and whose developmental function is enigmatic. We provide first experimental evidences that the ventrally located part of the CF has two distinct functions at two different phases of embryo development: (i) it works as a barrier at the onset of gastrulation by reducing forward cell flow and contributing to isolate the head cells, and (ii) it works as a tissue reservoir that is deployed via unfolding 40 minutes into gastrulation, driving backward cell flow that accompanies the extension of the germband during the so called slow phase^46,47^. The CF on the more ventral zone of the embryo is around 50 μm deep. This results in a non-negligible tissue reservoir of 100 μm in length along the AP axis (i.e., 20% of the *Drosophila* embryo length). While during the germband elongation fast phase (preceding the slow phase) the posterior midgut movement rules extension^27^, the unfolding of the CF could potentially have a net contribution to the extension of the germband during the slow phase.

We show that the mechanical forces initiating the formation of the CF are intrinsic to the CF region and are under the control of the RhoGEF2 and Dp114RhoGEF pathways that are also key during germband furrowing and extension^29,48^. We also show that ROCK dependent phosphorylation of MyoII is necessary to drive CF formation, supporting the idea that actomyosin contractility is a major player in this process. When ectopically activating MyoII in one single cell, we can qualitatively mimic the onset of CF formation characterized by the shortening of an IC and the formation of a groove, but we cannot reproduce tissue involution. This holds true also when ectopically activating MyoII in an IC, demonstrating that CF formation starts with but is not triggered by IC apical-basal shortening. A higher-order timed molecular program is potentially at play setting the stage (e.g., by converting the blastoderm tissue from a non-excitable to an excitable cellular medium) and eventually providing the kick-off. The CF and the ventral furrow onset are simultaneous even if the two folds are located at different embryo positions and are under the direct control of different signaling factors. Together they both mark the onset of gastrulation and therefore the blastula-to-gastrula transition. Future work is necessary to unveil the mechanisms and their temporal control responsible for gastrulation kick-off.

During the formation of the CF, MyoII is present on the apical, lateral and basal sides of ICs. We show that contractility on the lateral side of cells is more effective to drive apical-basal shortening and groove formation than apical and basal contractility that have lesser and no shortening contribution, respectively. While lateral contractility drives apical-basal cell shortening, basal acute downregulation of MyoII and basal tension is necessary and sufficient to induce cell rolling over. The CF is a narrow fold formed by the involution of cells from both the head and trunk region. This results in medial apices of originally non-neighboring cells coming in close proximity. While junctions are known to form between lateral sides of epithelial cells, here we show that *de novo* adherens junctions form also between cell medial apices. These medial junctions sustain and could lock the CF in place, preventing it from precocious unfolding during the development of the embryo. 40 minutes into gastrulation the ventral part of the CF unfolds, driving backward cell flow. Future work is necessary to understand what drives unfolding. Medial apex-to-apex adhesion between head and trunk cells could be downregulated to unlock the CF in a timely fashion.

The CF is initiated on the left and right side of the embryo by a small group of cells that we dubbed first initiator cells (FICs) and it propagates towards the embryo ventral and dorsal side mirrored by a MyoII propagation wave. On the ventral side, the CF eventually intersects the ventral furrow. On the dorsal side, the left and right folds of the CF merge into each other, faithfully following the most anterior ‘stripe’ of the Eve transcription factor^14^ (part of the AP gene patterning system) that forms a continuous ring like pattern around the embryo^48^. How is the DV position of FICs controlled? While the AP gene patterning provides a signal modulation (i.e., cell positional information) along the AP axis^11^, in principle, it would be per se insufficient to provide instructive information along the DV axis. We show that the DV patterning system controls the DV FIC position. In addition, both AP and DV signaling are necessary to ensure the CF MyoII wave. Therefore, the CF is positioned and dynamically shaped under the cross-control of AP and DV patterning genes. This provides another striking example of how the AP and DV molecular pathways can work together to provide a multidimensional instructive signal controlling the shaping of a tissue during morphogenesis^48^. By using combined multi-view light sheet microscopy, IR fs laser, two-photon optogenetics and image processing techniques, we show that the FICs function as a pacemaker, initiating the propagation of a mechanical trigger wave that sculpt the CF. For example, when disrupting the FICs on only on the left side of the embryo, the CF left fold is inhibited while the right fold is preserved. Eventually, the right fold propagates across the dorsal side towards the left side, rescuing the left dorsal-lateral part of the CF. In contrast, the right fold does not propagate across the ventral side and so the left ventral-lateral part of the CF will not be rescued. This shows that the formation of the dorsal side of the CF is more robust than the ventral side and it supports the idea that the ventral furrow functions as a barrier ventrally decoupling the right from the left fold of the CF. During CF propagation, groups of cells in the head region close to the CF (e.g., cells belonging to the first mitotic domain) undergo synchronous AP oriented division^49^. In addition, the posterior midgut moves anteriorly towards the dorsal head domain. These cellular and morphogenetic processes could eventually contribute to the folding of the dorsal part of the CF.

Similarly to the CF, the ventral furrow also propagates bi-directionally from a medial point. Nevertheless, while CF propagation is driven by a trigger wave, ventral fold propagation results from long-range tissue interactions in the absence of contractile waves^7^. A contractile wave has been reported also during the movement of the posterior midgut^50^. It is not yet determined if this wave plays a role in posterior morphogenetic movements. MyoII trigger waves could result from the transduction of a mechanical stimulus into a molecular signal^51-53^ between adhering cells. The induced signal could in turn drive a new mechanical stimulus in a recursive fashion across a tissue^54,55^. In the CF region junctions are relocated apically^31^. Apical junctional relocation could be driven by apical MyoII recruitment (preceding IC lateral MyoII recruitment) as, for instance, reported during ventral furrow formation^56^. Focusing of junctions at the apical side could reinforce the mechanical coupling between CF cells and eventually mediate the propagation of a mechanically transduced signal across the tissue. Based on this idea, two families of CF cells can be identified: the ICs that would change shape in response to a mechanical stimulus and the FICs that act as a pacemaker and which cell shape change would be mechano-transduction independent and under the control of a higher order timed genetic program. What is the advantage of forming a furrow via a propagating mechanism? A mechanical trigger wave could be an energetically efficient and robust morphogenetic mode to insure a spatio-temporally continuous tissue shape change resulting in a smooth and coherent structure or organ. Future work is necessary to better understand the molecular machinery underlying the propagation of mechanical waves in epithelial tissues. This knowledge could be used in the near future to synthetically build and shape functional organs^57^.

## Acknowledgments

We thank S. De Renzis, E. Wieschaus, S. Streichen, Y. Bellaiche, T. Lecuit and C. Collinet for providing fly stocks and compounds; B. Delorme for developing ImageJ tools for image analysis and for helping with 3D cell segmentation; Stéphane Noselli, Matej Krajnc and all members of the Rauzi lab for critical reading of the manuscript and fruitful discussions; the PRISM imaging facility for technical support. M.R. thanks the company Luxendo Bruker and SPARK LASERS for fruitful collaborations and support. This work was supported by the French government through the UCA^JEDI^ Investments for the Future project managed by the National Research Agency (ANR-15-IDEX-01), the Investments for the Future LABEX SIGNALIFE (ANR-11-LABX-0028-01), the Tramplin-ERC program of the National Research Agency (ANR-16-TERC-0018-01), the ATIP-Avenir program of the CNRS; the Human Frontier Science Program (CDA00027/2017-C); the Region SUD PACA Research and ERC Booster programs.

## Author Contributions

MR designed the project together with AP. AP and MR planned the experiments. AP performed the experiments. MR planned and SP performed the electron microscope trials. AP performed data processing and quantifications. AP and MR analyzed the data. MR wrote the manuscript together with AP.

