## Supplementary Figures for "A mechanical wave travels along a genetic guide to drive the formation of an epithelial furrow"

#### SUPPLEMENTARY FIGURE LEGEND

**Supplementary Figure 1. CF initiation is driven by intrinsic forces.**
**a** CF depth as a function of time for a wildtype and a cauterized embryo. **b** En-face view (left) and orthogonal view (right) along the dashed white line for a wild type and a cauterized embryo (top and bottom, respectively). t = 0 is the beginning of apical narrowing of the IC (magenta). The laser cauterization position is indicated by the red dashed line and arrowheads. Scale bar 30 µm.

**Supplementary Figure 2. RhoGEFs play a role in sculpting the CF.**
**a** Top panel: representative time-lapse images showing confocal en face views of control embryo, expressing membrane marker and Eve protein (purple) and the corresponding cell surface area over time. Blue indicates smaller cell surface area. t = 0 corresponds to the end of cellularization. Bottom panel: normalized IC cell surface area in different genetic backgrounds. **b** Delay of CF left and right fold initiation. **c** MyoII activation of a cell outside the CF region. The red rectangle denotes the photo-activated region. The red arrowhead indicates the ectopic fold (EF) position. White arrowhead indicates the CF position. Scale bar 20 µm.

**Supplementary Figure 3. Apical, lateral and basal contraction during tissue folding.**
**a, b, c** MyoII subcellular activation on the apical (**a**), lateral (**b**) and basal (**c**) sides. Apical displacement is measured as the distance between the vitelline membrane and the apical side of photo-activated cells. Error bars indicate ±SD. For apical activation, n = 8 cells from 3 embryos. For lateral activation, n = 7 cells from 3 embryos. For basal activation, n = 4 cell from 2 embryos. **d** Single junction ectopic MyoII activation. The red rectangle denotes the photo-activated region. Scale bar 30 µm (**a**, **b**, **c**), 15 µm (**d**).

**Supplementary Figure 4. Basal contractions and cell-cell adhesion during CF internalization.**
**a**. IR fs laser ablation of the basal actomyosin network. Upper panels: apical surface views, membrane (red) and MyoII (green). Middle panels: basal views. Bottom panels: orthogonal views. t = 0 is the first time point after ablation. Scale bar 30 µm. **b**. E-cad and membrane distribution in the CF. The arrowhead highlights E-cadherin clusters between the apices of opposite (head and trunk) cells in the CF. Scale bar 20 µm.

**Supplementary Figure 5. CF MyoII waves are under the control of AP and DV gene patterns.**
**a, b, c**. Embryo cylindrical projections showing the distribution of MyoII during CF formation in a wild type embryo (**a**), an embryo from a *dorsal*- female (**b**), and an embryo from a *bnt*- female (**c**). Dorsal (D), ventral (V), lateral left (LL) and lateral right (LR). t = 0 corresponds to the end of cellularization (cell length, 30 µm). Arrows show the spreading of the MyoII wave. Scale bar 100 µm. **d**. Time alignment of different morphogenetic processes during gastrulation in wild type embryos, embryos from *dorsal*- females and *dpp*- embryos.

**a**

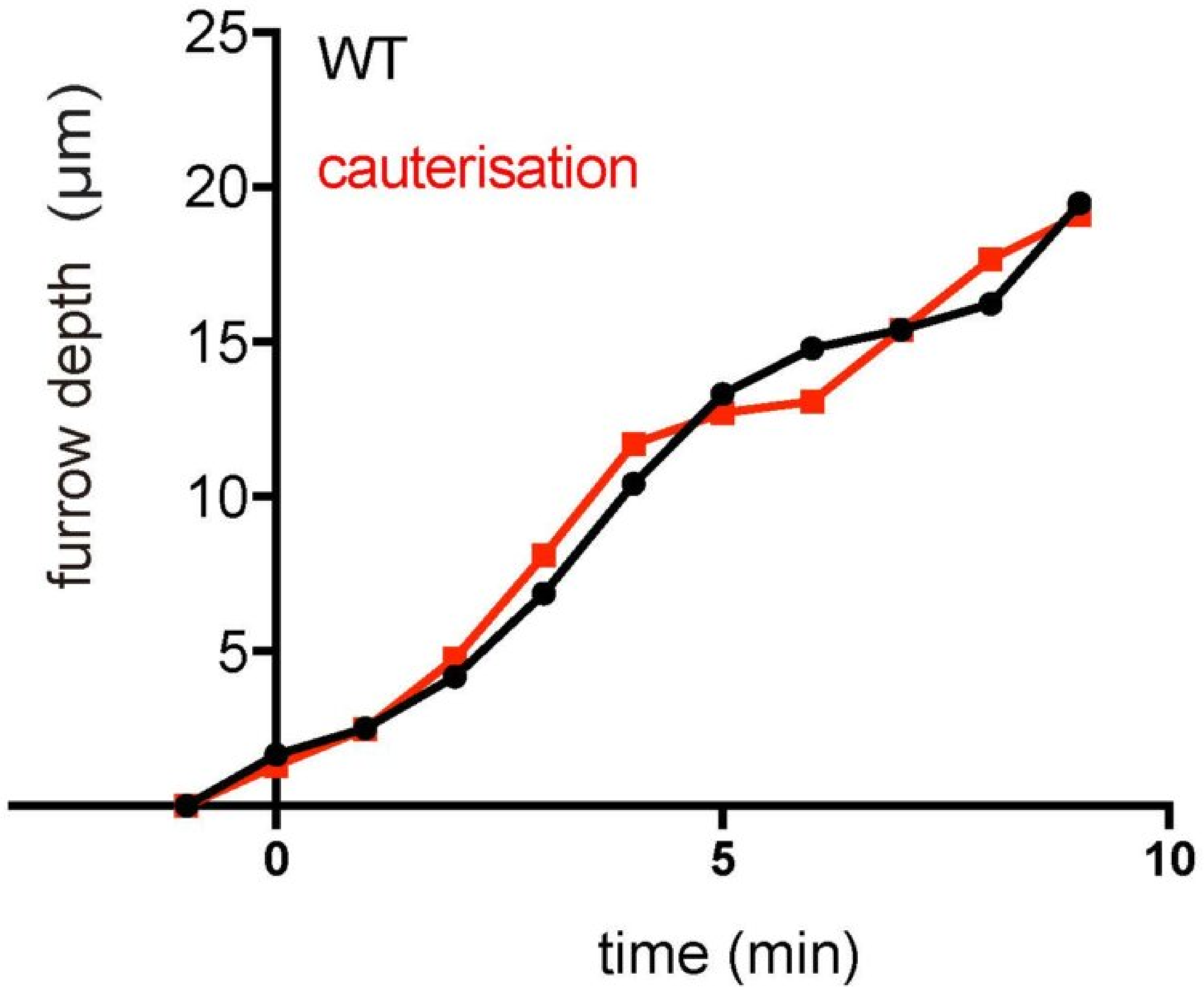


**b**

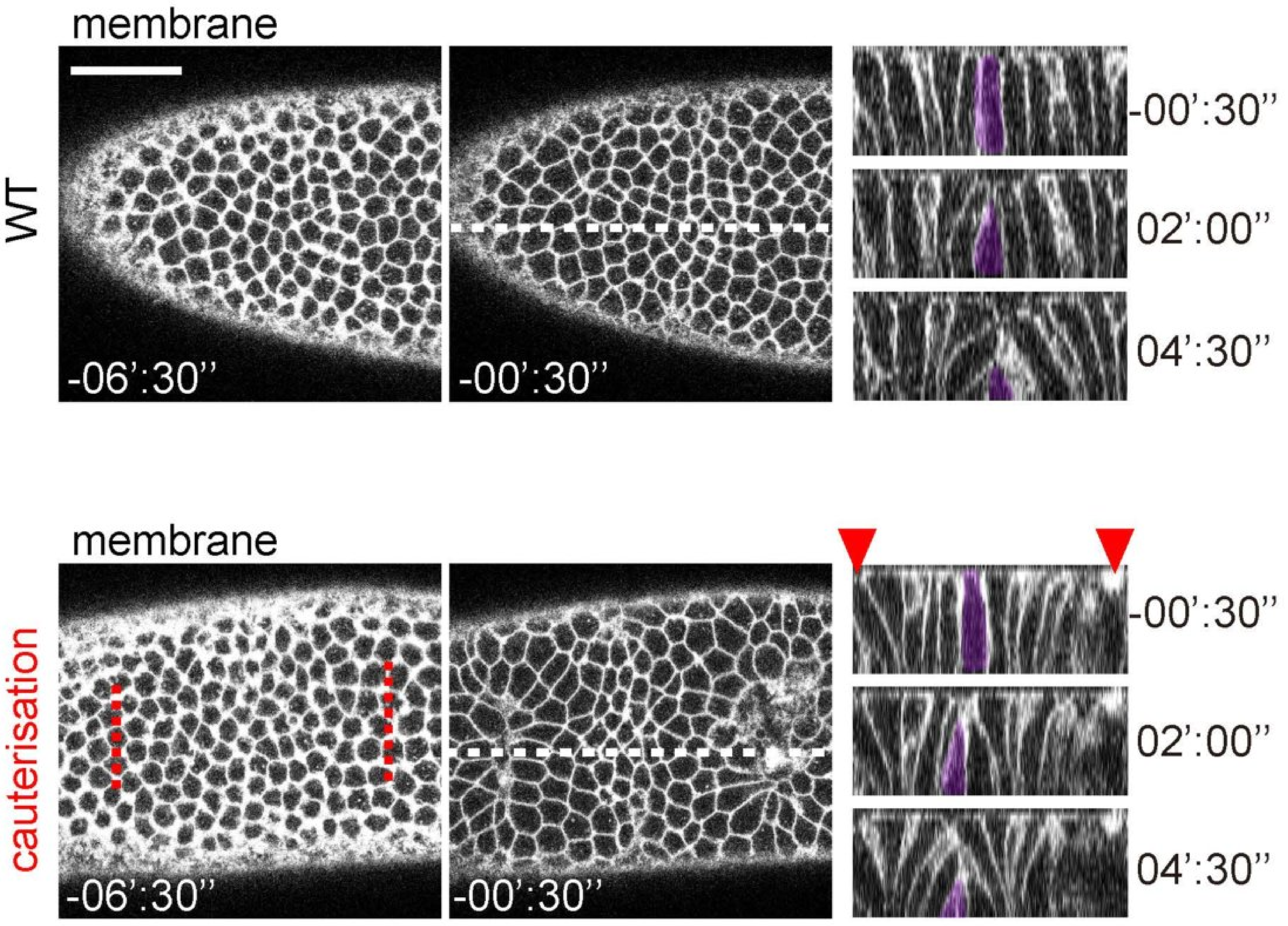


**Supplementary Figure 1**

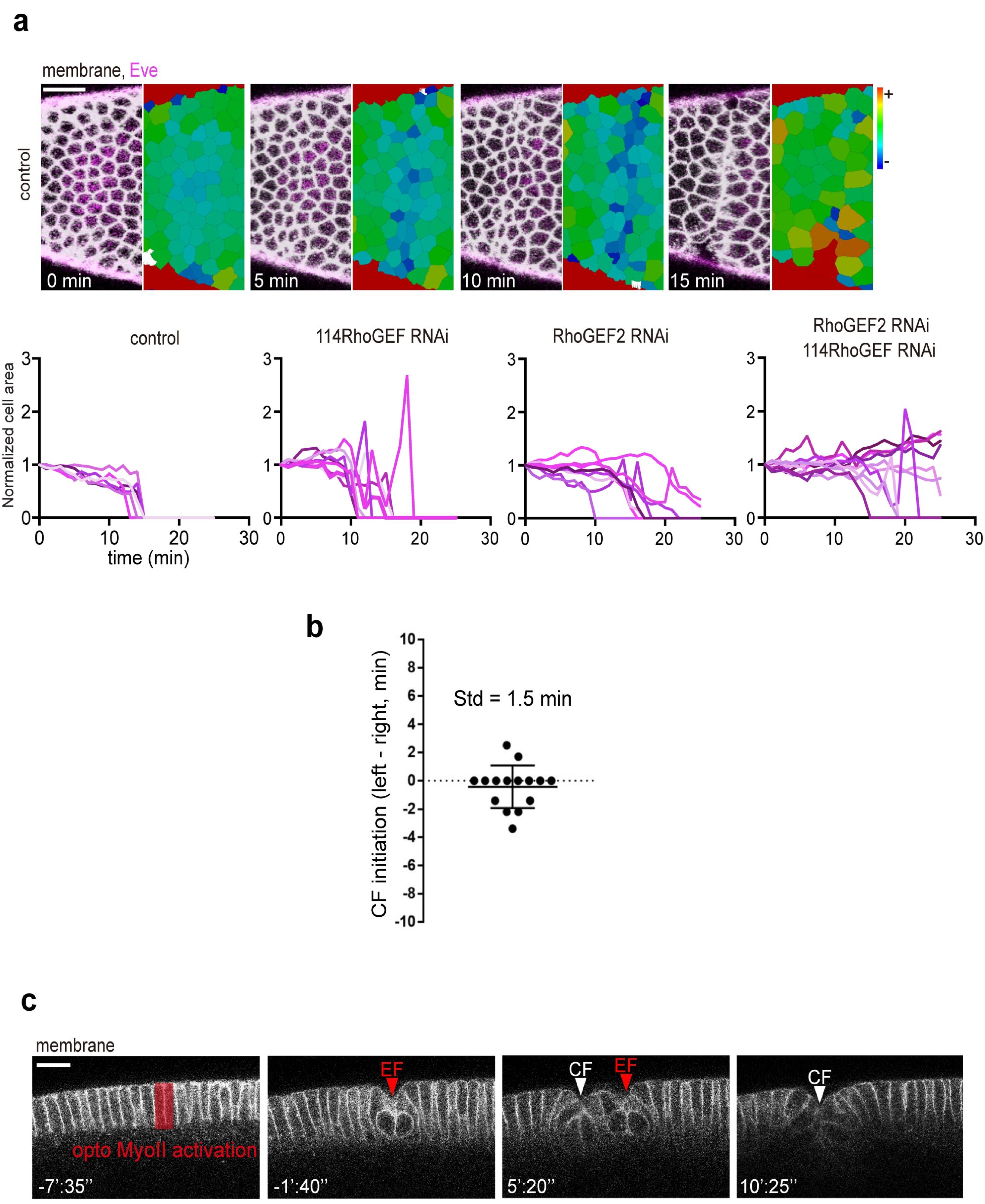


### Supplementary Figure 2

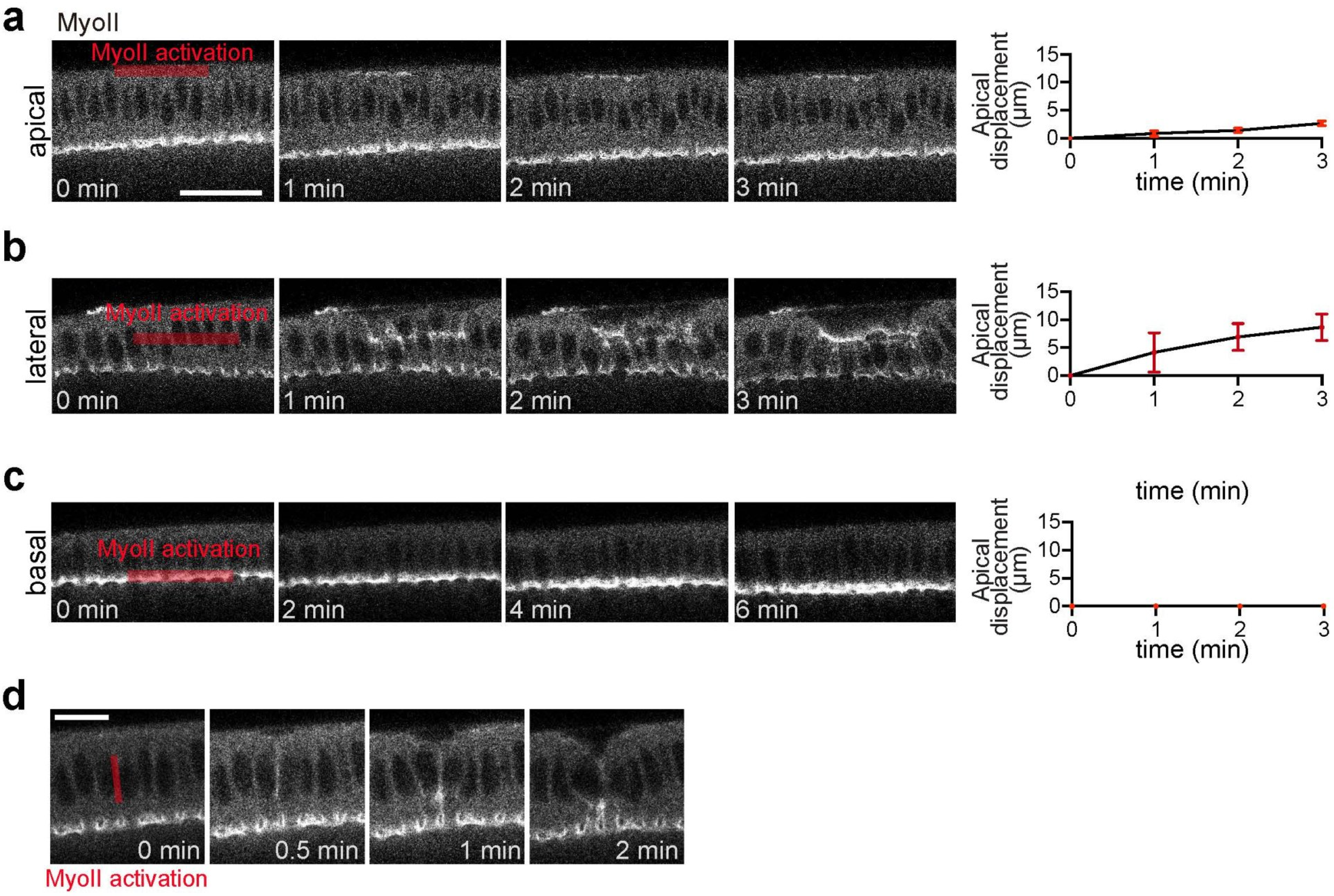


**Supplementary Figure 3**

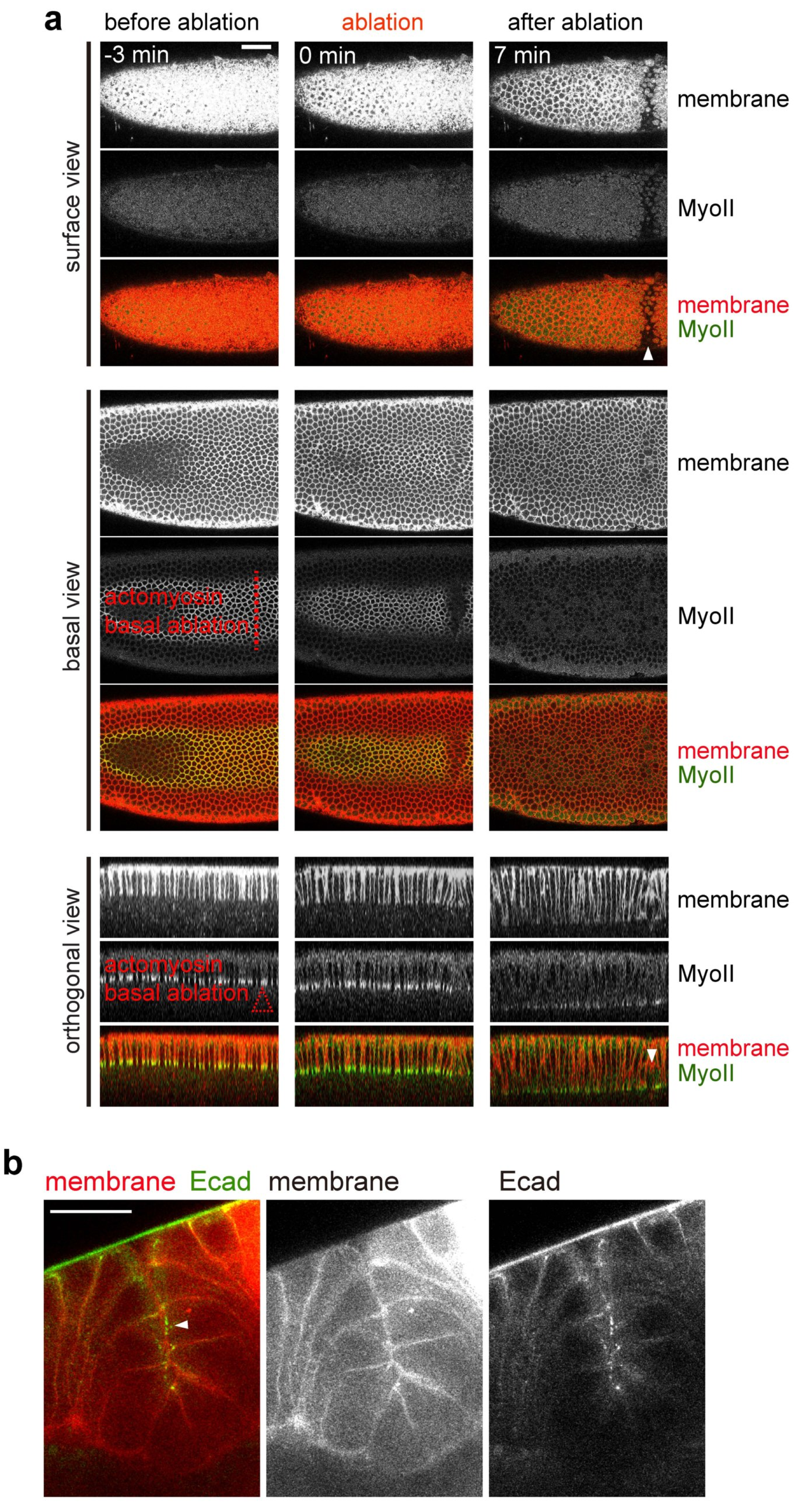


### Supplementary Figure 4

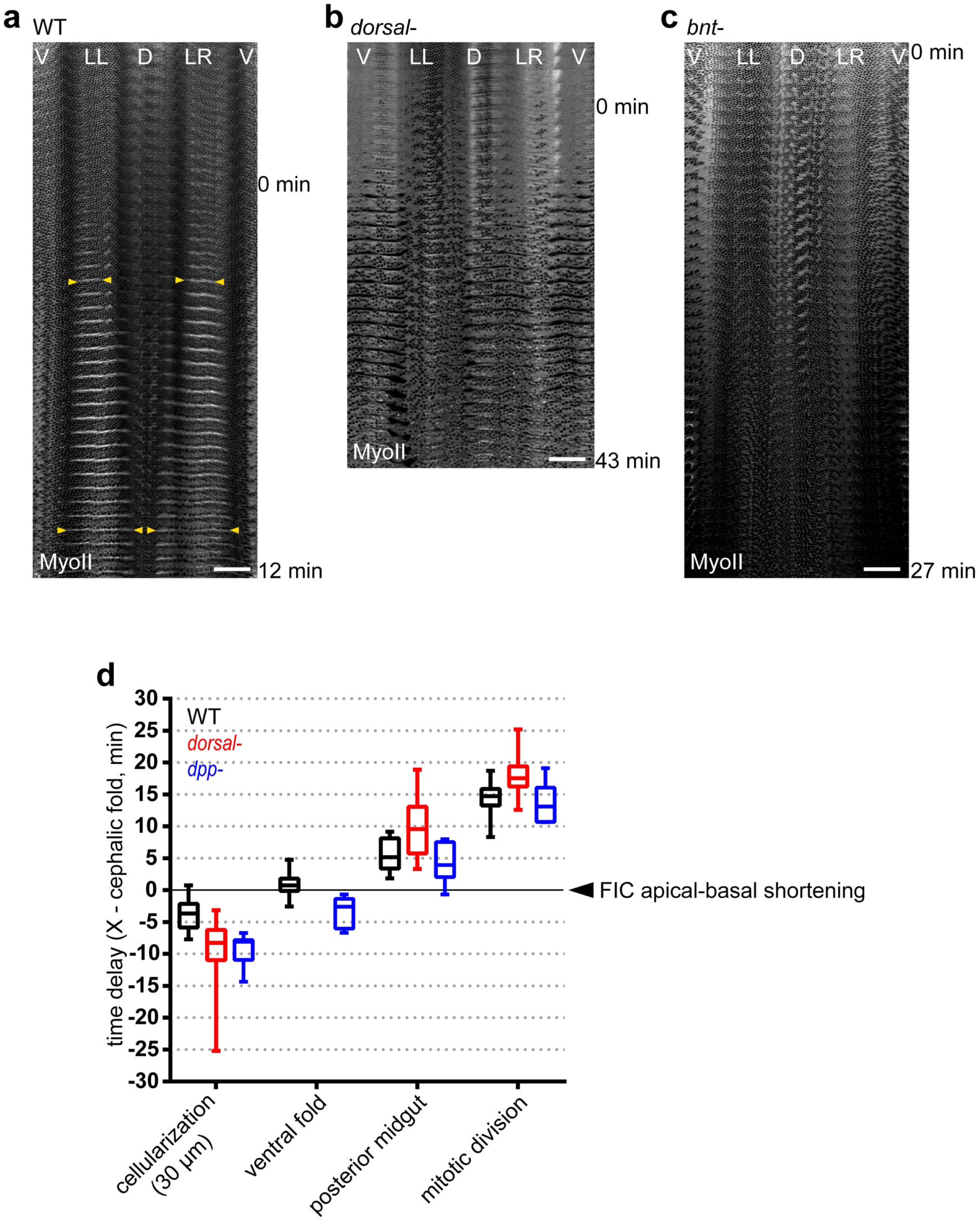


### Supplementary Figure 5
