## Supplementary Information for "A mechanical wave travels along a genetic guide to drive the formation of an epithelial furrow"

### METHODS

#### *Drosophila* stocks and genetics

*Drosophila melanogaster* stocks used in this study and associated references are listed in Supplementary Table 1. All stocks and crosses were maintained at room temperature. Fly stock expressing mCherry fusion to Gap43 was used for membrane labeling. MyoII localization was observed using the fly stocks expressing mCherry or GFP fusion to *Drosophila* Myosin regulatory light chain (MRLC), Spaghetty squash (Sqh), under the control of *sqh* promoter. Even-skipped (Eve) protein expression was monitored using the fly stock Eve::YFP. To monitor cell dynamics during CF formation (Fig. 1a, 1d and Fig. 4a, 4b), embryos from *klarsicht* (*klar-*) homozygous mothers expressing Gap43::mCherry were used. For cauterization, embryos were obtained from crosses of females expressing Gap43::mCherry or Sqh::mCherry with males from the stock Eve::YFP. To achieve RhoGEF2 and Dp114RhoGEF silencing, embryos were collected from crosses of females $_{p}$*ubi: Gap43::mCherry/+; tub*$^{mat}$ *Gal4/+; UAS RhoGEF2 RNAi/tub*$^{mat}$ *Gal4* or $_{p}$*ubi: Gap43::mCherry/+; UAS Dp114RhoGEF RNAi/ tub*$^{mat}$ *Gal4; tub*$^{mat}$ *Gal4/+* or $_{p}$*ubi: Gap43::mCherry/+; UAS Dp114RhoGEF RNAi/ tub*$^{mat}$ *Gal4; UAS RhoGEF2 RNAi/t tub*$^{mat}$ *Gal4* with males from the stock Eve::YFP. For the photorecruitment of RhoDN experiment, embryos were collected from *UASp CIBN::pmGFP/ tub*$^{mat}$ *Gal4; UASp Cry2::RhoDN::mCherry/ tub*$^{mat}$ *Gal4* mothers. For photorecruitment of RhoGEF2, embryo were collected from $_{p}$*ubi: Gap43::mCherry/+; UASp CIBN::pmGFP/ +; UASp Cry2::RhoGEF2/*$_{p}$*osk Gal4* or *UASp CIBN::pmGFP/ Sqh::mCherry; UASp Cry2::RhoGEF2/*$_{p}$*osk Gal4* mothers. For basal photorecruitment of RhoGEF2, embryos were collected from *UASp PatJ::CIBN::pmGFP/Sqh::mCherry; UASp Cry2::RhoGEF2/*$_{p}$*osk Gal4* mothers. For light-mediated actin depletion, embryos were collected from *UASp CIBN::pmGFP/ tub*$^{mat}$ *Gal4; UASp Cry2::dOCRL/ tub*$^{mat}$ *Gal4* or $_{p}$*ubi: Gap43::mCherry/+; UASp CIBN::pmGFP/ tub*$^{mat}$ *Gal4; UASp Cry2::dOCRL/ tub*$^{mat}$ *Gal4* mothers. Maternal mutant embryos for the gene *dorsal* (*dl*) were collected from cross of females $_{p}$*ubi: Gap43::mCherry/+; dl[1]/dl[1]* with males from the stock Eve::YFP or from females *dl[1]/dl[1]; Sqh::eGFP*. Zygotic mutant embryos for the gene *decapentaplegic* (*dpp*) were obtained from the cross $_{p}$*ubi: Gap43::mCherry/+; dpp[d6]/CyO x* $_{p}$*ubi: Gap43::mCherry; dpp[d6]/CyO*.

#### Time-lapse imaging and photomanipulation

Cages were maintained at 22 °C. Flies of the desired genotype were collected in cages with agar and yeast paste. Embryos were dechorionated in bleach, selected for appropriate stage, using a standard stereomicroscope under transmitted illumination, and mounted on glass-bottom plate in water. For light sensitive lines, cages were kept and manipulated in the dark. Additionally, the microscope light source was replaced with a conventional red-emitting LED lamp to prevent unwanted photoactivation. Time-lapse imaging of the mid-coronal section was performed with a Zeiss 880 inverted confocal microscope, Fast Airyscan option, with a 40×, numerical aperture 1.1 water-immersion lens (Zeiss LD-C Apo 421867-9970), with a 488 nm argon laser and a 561 nm DPSS laser. Imaging data was obtained using the Zeiss ZEN software. For drug injection, embryos were imaged on Nikon spinning disc microscope with 20x numerical aperture 0.75 objective (NIKON CFI Planfluor) or 40x numerical aperture 1.25 objective (NIKON CFI APO lambdaS), water-immersion, using 488 nm and 561 nm lasers. Data was acquired using Metamorph software.

For photoactivation experiments, live imaging was performed using a 780 NLO (Zeiss) inverted confocal microscope, equipped with the Spectra-Physics Mai Tai DeepSee IR fs laser (Newport Corp., Irvine, CA, UAS), with a 40×, numerical aperture 1.2 water-immersion lens (ZeissC-Apo 421767-9970), with a 488 nm argon laser and a 561 nm DPSS laser. Imaging data was obtained using the Zeiss ZEN software. Cry2 photorecruitment was obtained using the FRAP function of ZEN software. Photorecruitment of RhoDN was achieved on the mid-coronal section (around 50 µm from the dorsal side) by rectangular activation of the cortex using 950 nm, 32 mW laser power at the focal point, 50 iterations, 2.4 µs pixel dwell, over a z-stack. Photorecruitment of RhoGEF2 was achieved on the mid-coronal section by rectangular activation of the cortex using 950 nm, 18 mW laser power at the focal point, 30 iterations, 2.4 µs pixel dwell.

For light-mediated actin depletion combined with 3D light-sheet in toto imaging, a 920 nm IR femtosecond laser (Alcor, Spark Lasers), working at 88 fs pulse duration and 79.73 MHz repetition rate, was coupled with multi-view light sheet imaging (MuVi SPIM), as described in (de Medeiros et al 2020). Cry2-OCRL photorecruitment was carried out by scanning the region of interest – typically every 1-2 minutes for a maximum of 15 minutes – until clear Cry2 colocalization to the membrane was observed. After that, *in toto* imaging was maintained.

**In toto 4D imaging, digital reconstruction and data processing**

Embryos were dechorionated in bleach, selected for the appropriate stage and mounted in a glass capillary filled with 0.5% gelrite, with their AP axis parallel to the capillary. A small portion of the gelrite cylinder containing the embryo was pushed out to image on Luxendo MuVi SPIM with an Olympus 20x 1.0 NA objective, using 488 nm and 594 nm lasers. Z-stacks were acquired with a step-size of 1 µm, during each acquisition embryos were imaged from two opposing directions simultaneously and successively from two directions orthogonal to the first two (0° = dorsal and ventral; 90° = lateral view). Thus, for every single time point, four 3D stacks of the embryo were recorded. The four stacks at each time point were fused resulting in a combined single image with isotropic pixel resolution. The 4D image analysis code ASTEC (Guignard et al 2020) was used to fuse and align the stacks. The unrolling was performed to extract the apical surface of the embryo following the Matlab protocol described in (Rauzi et al 2015).

**Laser manipulations**

For laser manipulations in WT embryos, live imaging was performed using a 780 NLO (Zeiss) inverted confocal microscope, equipped with the Spectra-Physics Mai Tai DeepSee IR fs laser (Newport Corp., Irvine, CA, UAS), with a 40×, numerical aperture 1.2 water-immersion lens (ZeissC-Apo 421767-9970), with a 561 nm DPSS laser. Actomyosin network ablations were obtained by exposing the network to the focused beam with an average power of 140 mW (at 950 nm with 90% transmission) on the focal plane, using the FRAP function of the ZEN software (4 iterations, pixel dwell 3.6 µs). For cauterization, the IR laser was tuned to 930 nm with 80% transmission, 2 iterations, pixel dwell 50 µs. For vitelline membrane perforations, the IR laser was tuned to 840 nm with 100% transmission, 4 iterations, pixel dwell 3.6 µs.

**Drug injections**
Embryos were dechorionated in bleach, selected for the appropriate stage, dried in a box with silica beads for 9 minutes and mounted in halocarbon oil before injection. To inhibit MyoII activity, 50 mM Rok kinase inhibitor, Y-27632 (TOCRIS), was injected into embryos at mid-cellularization stage. Water injected embryos were used as a control.

**RNA interference against shotgun (E-cad)**
The dsRNA probe against *shortgun* (E-cad) was generated by PCR using the following primers, where the sequence in small letters is T7 promoter and the sequence in capital letters corresponds to the template sequence.
shg2-T7-Fp taatacgactcactatagggcGAGTCTGTTTGATAATGGTGAGC
shg2-T7-R taatacgactcactatagggcGGTTTCCATCGTTCTGGTGAATC
Embryos obtained from 30 min egg collection were dried, injected with 5 μM dsRNA and imaged 2hrs after injection using 780 NLO (Zeiss) inverted confocal microscope.

**Electron microscopy**
Drosophila embryos of different stages were dechorionated in bleach and then cryo-immobilized by High Pressure Freezing (HPF) in a Ficoll paste (Leica EM ICE) followed by freeze substitution in the Leica EM AFS 2 in the medium precooled to -90°C (0.1 % tannic acid, 1% OsO4 and 0.1 % uranyl acetate in acetone). At the end of the process (4 days), when the samples reach the room temperature they are washed with acetone several time and then substituted in acetone-Epon mixture, then infiltrated in EPON and polymerised at 60°c overnight. Ultrathin sections (80 nm) were contrasted with uranyl acetate (4% in water) then lead citrate and viewed in a Transmission Electron Microscope (JEOL JEM 1400) operating at 100 kV and equipped with an Olympus SIS MORADA camera.

**Vertex displacement analysis**
Homozygous *klar* mutant embryos expressing Gap43::mCherry were imaged on Zeiss 880 confocal microscope. Time lapse images of mid-coronal section were aligned temporally based on initiation - the descent of cell apex of cell identified as the IC. Apical side of each vertex was manually tracked using Fiji.

**4D cell segmentation and rendering**
ASTEC (Guignard et al 2020) was used for 4D segmentation and IC volume analysis. Imaris was used for 3D rendering.

**RhoGEF RNAi analysis**
Time-lapse en face images of embryos expressing either Gap43::mCherry and Eve::YFP along or with one or two UAS RNAi constructs were obtained on a 780 NLO (Zeiss) inverted confocal microscope. Movies were aligned based on the timing of the end of cellularization, defined by the appearance of the membrane signal in the CF region (identified by Eve expression) 30 μm from the apical surface.

**Furrow depth measurement**
Furrow depth was measured, using Fiji, as the distance between the vitelline membrane and the apex of either the photoactivated cell (for etopic MyoII activation) or the endogenous IC, identified by the descent of the cell apex.

**MyoII intensity measurement**

MyoII intensities were measured in the cytoplasm and cortex, using Fiji, and the ratio between cortical to cytoplasmic MyoII was calculated.

**FIC position measurement**

The FIC position was measured on unrolling views, using Fiji, as the distance between the FICs and the ventral midline, and normalized by the distance between the dorsal and ventral midlines.

**2D cell segmentation and tracking**

For the RNAi analysis RhoGEFs and experiments testing CF barrier function, 2D cell segmentation and tracking were performed using IMARIS.

**Statistics**

The Mann-Whitney test (GraphPad Prism software) was used to make statistical comparisons of results among groups. A value of $P<0.05$ was considered to be statistically significant while a value of $P<0.001$ was considered to be remarkably statistically significant.

### MOVIE LEGENDS

**Movie 1**. Time-laps movie of surface view in a membrane-labeled embryo starting at a late cellularization stage. Top: control embryo. Bottom: immobile boundaries generated along the DV axis around the furrowing zone before the onset of CF initiation. t = 0 corresponds to IC apical narrowing. Scale bar 30 µm.

**Movie 2**. 3D embryo rendering of the CF region in a membrane-labeled *klar*- embryo. Cell segmentation was performed from 3D time-lapse *in toto* imaging. IC, magenta. t = 0 is the onset of CF formation. Scale bar 30 µm.

**Movie 3**. Time-lapse movie of surface view of a membrane-labeled embryo starting at the mid-cellularization stage. Top: control embryo injected with water. Bottom: embryo injected with Roh kinase inhibitor (Y-27632). t = 0, 30 µm cellularization depth (onset of CF formation).

**Movie 4**. Top: Time-lapse movie of mid-coronal view in a membrane-labeled embryo at mid-cellularization stage expressing CIBN::pmGFP and Cry2::RhoGEF2, with a photo-activated cell outside of the CF region. Red rectangle denotes the photo-activated cell. t = 0 corresponds to the endogenous CF initiation. Scale bar 15 µm.

**Movie 5**. Top: Time-lapse movie of mid-coronal view in a membrane-labeled embryo at mid-cellularization stage expressing CIBN::pmGFP and Cry2::RhoGEF2, with a photo-activated presumptive IC. Middle: zoom on photo-activated (left) CF region. Bottom: zoom on control (right) CF region. Red rectangle denotes the photo-activated cell. t = 0 corresponds to the endogenous CF initiation. Scale bar 30 µm.

**Movie 6**. Time-lapse movie of mid-coronal view in a MyoII-labeled embryo. t = 0 is the onset of CF formation. Scale bar 30 µm.

**Movie 7.** Time-lapse movie of mid-coronal view in a membrane-labeled embryo. IR fs laser perforation (arrow head) of the vitelline membrane in the proximity of the CF. Scale bar 25 µm.

**Movie 8.** Cylindrical project time-lapse movie of the CF region with MyoII-labeled of a wild type embryo. Dorsal is in the center while ventral is both on the right and left sides (see ventral furrow formation). Scale bar 100 µm.

**Movie 9.** Cylindrical project time-lapse movie of the CF region with membrane-labeled of a wild type embryo. Ventral (V), lateral left (LL), dorsal (D), lateral right (LR). Scale bar 100 µm.

**Movie 10.** Cylindrical project time-lapse movie of the CF region with membrane-labeled of a *dpp*- embryo. Ventral (V), lateral left (LL), dorsal (D), lateral right (LR). Scale bar 100 µm.

**Movie 11.** Cylindrical project time-lapse movie of the CF region with membrane-labeled of an embryo laid by *dorsal*- female. Scale bar 100 µm.

**Movie 12.** Cylindrical project time-lapse movie of the CF region with MyoII-labeled of an embryo laid by *dorsal*- female. Dorsal is in the center while ventral is both on the right and left sides (see ventral furrow formation). Scale bar 100µm.

**Movie 13.** Cylindrical project time-lapse movie of the CF region with MyoII-labeled of an embryo laid by *bnt*- female. Dorsal is in the center while ventral is both on the right and left sides (see ventral furrow formation). Scale bar 100 µm.

**Movie 14.** Cylindrical project time-lapse movie of the CF region labeled with membrane and Cry2-OCRL. The red rectangle denotes the region of two-photon activation on the lateral left side of the embryo. The yellow line denotes the part of the CF that is formed. Dorsal is in the center while ventral is both on the right and left sides (see ventral furrow formation). Scale bar 100 µm.

**Movie 15.** Cylindrical project time-lapse movie of the CF region labeled with membrane and with Cry2-OCRL. The red rectangle denotes the region of two-photon activation on the ventral-lateral left side of the embryo. The yellow line denotes the part of the CF that is formed. Dorsal is in the center while ventral is both on the right and left sides (see ventral furrow formation). Scale bar 100 µm.

**Movie 16.** Time-lapse movie of the CF unfolding in a control condition (i.e., with no photo-activation). The ventral side of the embryo is shown and Cry2-OCRL and membrane are labeled. On the right, cell trajectories: blue is forward movement (towards the anterior), red is backward movement (towards the posterior and green is no movement. Scale bar 50 µm.

**Movie 17.** Time-lapse movie of the CF unfolding in a bi-lateral activation condition (i.e., with photo-activation on both the left and right side). The ventral side of the embryo is shown and Cry2-OCRL and membrane are labeled. On the right, cell trajectories: blue is forward movement (towards the anterior), red is backward movement (towards the posterior) and green is no movement. Scale bar 50 µm.

**Movie 18.** Time-lapse movie of the CF unfolding in a single-lateral activation condition (i.e., with photo-activation on only one side, the left side of the embryo in this case). The ventral side of the embryo is shown and Cry2-OCRL and membrane are labeled. On the right, cell trajectories: blue is forward movement (towards the anterior), red is backward movement (towards the posterior) and green is no movement. Scale bar 50 µm.
